# Estimating allele-specific expression of SNVs from 10x Genomics Single-Cell RNA-Sequencing Data

**DOI:** 10.1101/2019.12.22.886119

**Authors:** N M Prashant, Hongyu Liu, Pavlos Bousounis, Liam Spurr, Nawaf Alomran, Helen Ibeawuchi, Justin Sein, Dacian Reece-Stremtan, Anelia Horvath

**Author notes:** Shared contribution (N M Prashant, Hongyu Liu).

## Abstract

With the recent advances in single-cell RNA-sequencing (scRNA-seq) technologies, estimation of allele expression from single cells is becoming increasingly reliable. Allele expression is both quantitative and dynamic and is an essential component of the genomic interactome. Here, we systematically estimate allele expression from heterozygous single nucleotide variant (SNV) loci using scRNA-seq data generated on the 10x Genomics platform. We include in the analysis 26,640 human adipose-derived mesenchymal stem cells (from three healthy donors), with an average sequencing reads over 120K/cell (more than 4 billion scRNA-seq reads total). High quality SNV calls assessed in our study contained approximately 15% exonic and >50% intronic loci. To analyze the allele expression, we estimate the expressed Variant Allele Fraction (VAF_RNA_) from SNV-aware alignments and analyze its variance and distribution (mono- and bi-allelic) at different cutoffs for required minimal number of sequencing reads. Our analysis shows that when assessing SNV loci covered by a minimum of 3 unique sequencing reads, over 50% of the heterozygous SNVs show bi-allelic expression, while at minimum of 10 reads, nearly 90% of the SNVs are bi-allelic. Consistent with single cell studies on RNA velocity and models of transcriptional burst kinetics, we observe a substantially higher rate of monoallelic expression among intronic SNVs, signifying the usefulness of scVAF_RNA_ to assess dynamic cellular processes. Our analysis demonstrates the feasibility of scVAF_RNA_ estimation from current scRNA-seq datasets and shows that the 3’-based library generation protocol of 10x Genomics scRNA-seq data can be highly informative in SNV-based analyses.

## 1. Introduction

In the last several years, single cell RNA-seq (scRNA-seq) has become an accessible platform for genomic studies [1–3]. By enabling cell-level transcriptome analyses, scRNA-seq brings a major advantage over the conventional averaged bulk RNA-seq: the ability to assess intracellular relationships between molecular features. With the recent advances in the scRNA-seq technologies, estimations of genetic variation from scRNA-seq data are becoming reliable [4, 5] and several studies have demonstrated their usefulness in addressing key biological and clinical questions [6–18].

Genetic variants are traditionally called from DNA and often analyzed and interpreted in their context genotypes (for diploid organisms, homo- or heterozygous). For expressed loci, genetic variation can be also assessed from RNA-seq data [19–22], by calculating the variant allele fraction, (VAF_RNA_= n_var_ / (n_var_+ n_ref_), where n_var_ and n_ref_, are the variant and reference read counts, respectively). VAF_RNA_ is an informative measure of genetic variation for several reasons. First, as compared to the categorical genotypes (DNA allele count of 0, 1 and 2), VAF_RNA_ is a continuous measure allowing for precise allele quantitation, which is important for sites where VAF_RNA_ functions as a continuous metric. These include loci exhibiting preferential expression of functional alleles, somatic mutations in cancer, and RNA-editing loci. Second, in contrast to the (static) genotypes, VAF_RNA_ is dynamic and reflects the actual allele content in the system at any particular time, which allows for the assessment of dynamic and progressive processes [23–25]. Importantly, through primarily reflecting genetic variation, VAF_RNA_ is an essential component of the genomic interactome and plays a major role in phenotype formation [26–30].

However, a systematic analysis of the feasibility of VAF_RNA_ estimations from 3’-based scRNA-seq libraries, and its usefulness to address biological questions has not yet been performed. One of the basic biological processes assessed through VAF_RNA_ is the prevalence of random monoallelic expression (RME) across the diploid mammalian genome. Several recent scRNA-seq studies have described widespread RME in both human and murine models [15–17]. Most of these studies analyzed scRNA-seq data generated on full-length transcript platforms from hundreds of cells.

Here, we demonstrate a pipeline to estimate VAF_RNA_ from scRNA-seq data obtained from 10x Genomics Chromium platform [31]. We have selected this platform due to its growing popularity along with: (1) high throughput (our analysis includes 26,640 cells obtained from three healthy donors), (2) high depth of sequencing (~150,000 sequencing reads per cell), and, (3) support for unique molecular identifiers (UMI) for removal of PCR-related sequencing bias. Because VAF_RNA_ is sensitive to allele-mapping bias [24,25], we use SNV-aware alignments where reads mapped ambiguously due to the variant nucleotide are removed [32]. From the SNV-aware alignments, we systematically assess the ability to estimate VAF_RNA_ using three different thresholds for minimum required number of unique sequencing reads (minR) - 3, 5 and 10. We compare outputs across thresholds and individuals, and outline lists of consistent observations. We also demonstrate an approach to assess RME, and compare the results from scRNA-seq generated on 10x Genomics Chromium with studies based on different platforms.

## 2. Materials and Methods

### 2.1. Data

We used publicly available scRNA-seq data from 26,640 human cells from three healthy donors (N8, N7 and N5); the scRNA-seq data was generated on 10x Genomics Chromium v2 platform [31]. The library preparation and sequencing are described in detail elsewhere [31]. Briefly, cells were partitioned using 10x Genomics Single Cell 3’ Chips, and barcodes to index cells (14 bp) as well as transcripts (10 bp UMI) were incorporated. The constructed libraries were sequenced on an Illumina NovaSeq 6000 System in a 2× 150 bp paired-end mode.

### 2.2. scRNA-seq data processing

The processing pipeline is shown on Figure 1. First, we extracted cell barcodes and UMIs using UMI-tools from the pooled (per donor) raw sequencing reads [33]. Next, we aligned the reads to the latest version of the human genome reference (GRCh38, Dec 2013) using STAR v.2.7.3.c [34] in 2- pass mode with transcript annotations from assembly GRCh38.79. We then called SNVs in the pooled alignments using GATK v.4.1.4.1 [20]. We selected heterozygous SNVs based on the presence of minimum of 50 high quality reads supporting both (reference and alternative) nucleotide in the pooled alignments. From those, we retained for further analysis heterozygous SNVs matching the following requirements: QUAL (Phred-scaled probability) > 100, MQ (mapping quality) > 60, QD (quality by depth) > 2, and FS (Fisher’s exact test estimated strand bias) = 0.000). In addition, after annotation (SeattleSeq v.13.00, dbSNP build 150) we removed SNV loci positioned in repetitive or intergenic regions. The SNV lists for each donor were then used as an input for a second, SNV-aware alignment using STAR, this time including the WASP-option [32, 34] for removal of reads mapped ambiguously due to the variant nucleotide. The SNV-aware alignments were dedupplicated keeping the reads with the highest mapping scores using the UMIs, and demultiplexed using the cell barcodes. The raw gene counts were estimated using featureCount [35], after which the gene counts were normalized and scaled using Seurat v.3.0 [36]. The gene counts were then used to remove cells with low quality data, defined as less than 3,000 detected genes and/or mitochondrial genes’ expression higher than 6% of the total gene expression. We estimated VAF_RNA_ on the individual alignments from the cells with high quality data using ReadCounts [26] with three different thresholds for minimum required number of reads (minR): minR = 10, minR = 5, and minR = 3.

**Figure 1.**
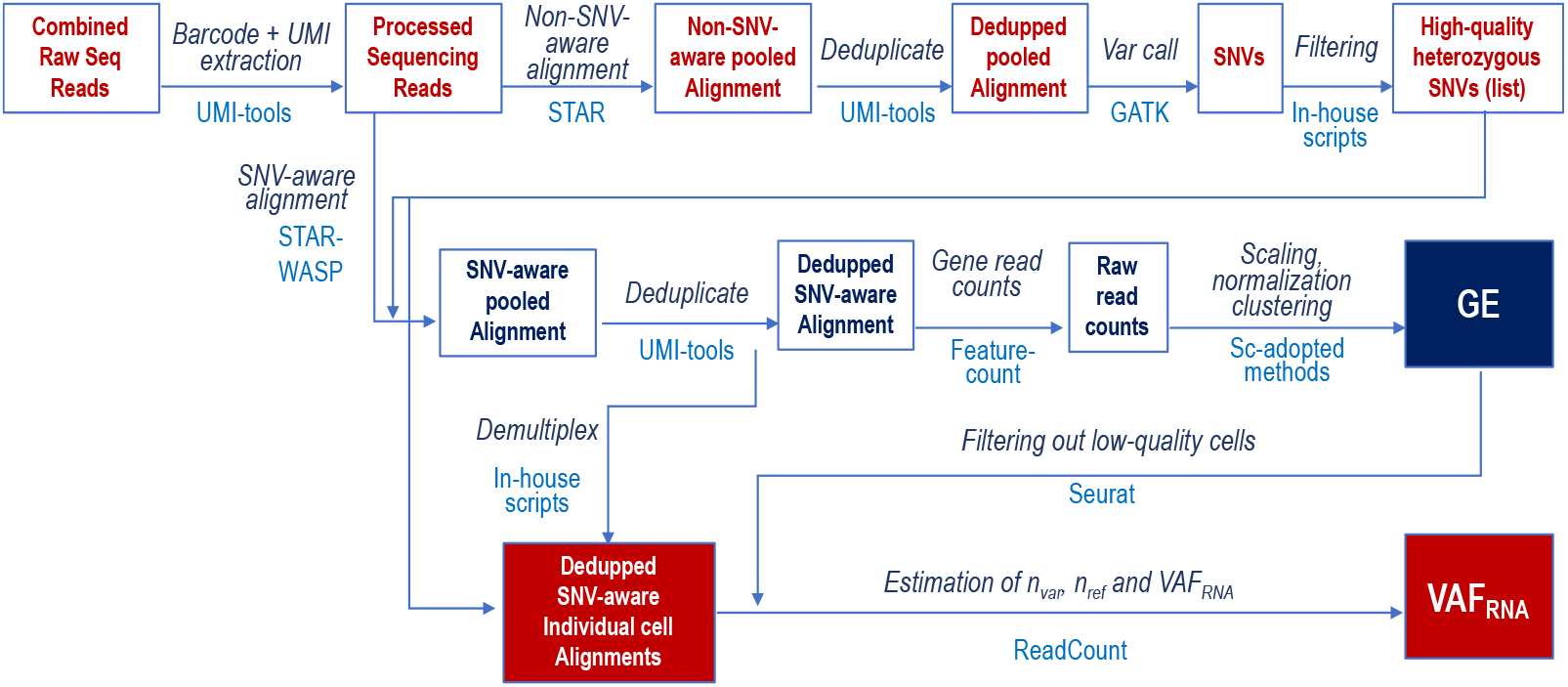
Analytical workflow for estimation of VAF_RNA_ from sc-RNA-seq data.

## 3. Results

### 3.1. Overall findings

The before- and after-filtering distributions of genes and sequencing reads is shown on Figure 2. The number of individual single cells with high quality data retained for further analysis was 8533, 8125 and 9115 for N8, N7 and N5, respectively. In these cells, we estimated VAF_RNA_ in 50,532 SNV loci in N8, 61,407 loci in N7, and 38,822 loci in N5, which were the number of loci retained after filtering for heterozygosity, quality, and position in intragenic non-repetitive regions. To support multi-cell estimations, we retained for statistical analyses only positions for which VAF_RNA_ is estimated from the selected number of unique sequencing reads (minR) from a minimum of 10 individual cells. Accordingly, unless otherwise indicated, the hereafter presented analyses are assessments from a minimum of 10 cells (per donor). For minR = 10, the absolute number of these positions was 366, 431 and 277 for N8, N7 and N5, respectively. This number was approximately 4-fold higher for positions assessed at minR = 5 and up to 20x-higher for positions at minR=3; the outputs are summarised in Table 1. We note that the relaxed thresholds are inclusive for the more stringent ones (i.e. minR = 5 loci include the loci at minR = 10, etc.). Of note, between 6 and 14% of all captured SNVs have been previously associated with clinical phenotype or highlighted by GWAS analyses (See Table 1).

**Figure 2.**
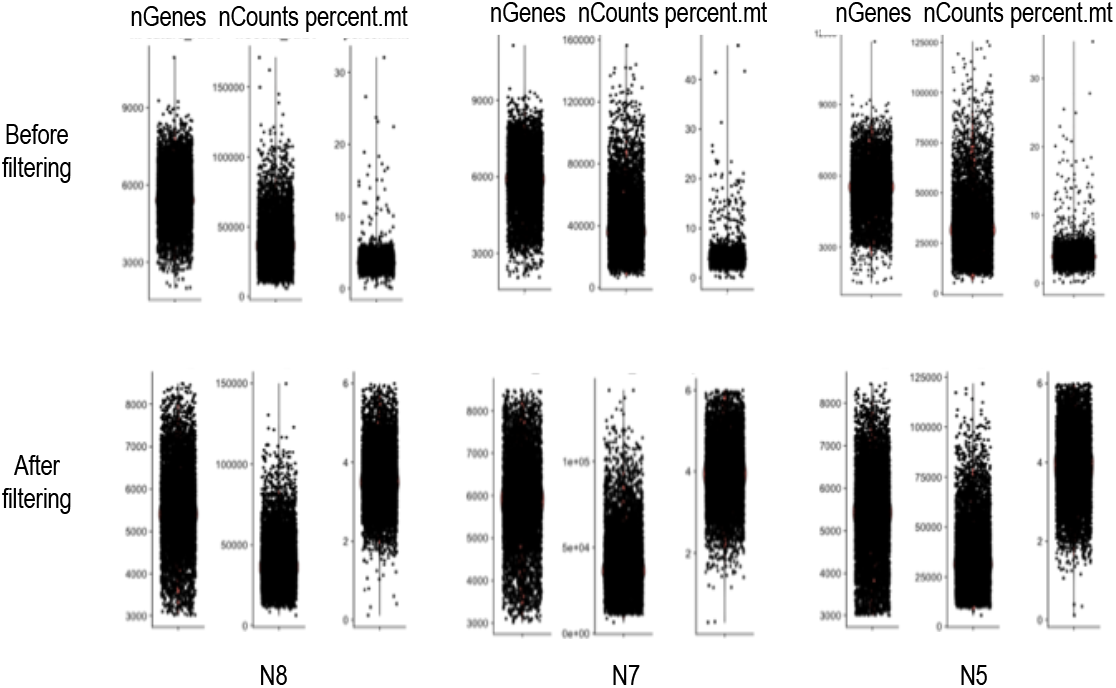
Number of genes, number of sequencing reads and percent mitochondrial genes for N8, N7 and N5 before (top) and after (bottom) filtering out of cells with low quality data.

**Table 1.** Summary statistics for the scRNA-seq data of the 3 ADSC samples. We estimated the number of SNV loci in at least 10 individual cells with different requirements for minimum number of unique reads (minR), and from those, the number of SNVs associated with phenotype.

| Sam<br>ple | N<br>cells | N<br>reads | Mean<br>reads/<br>cell | Median<br>genes/<br>cell | N cells<br>post<br>filtering | N Het SNVs<br>(min 10 cells) |  |  | N SNVs with<br>phenotype/clinics |  |  |
| --- | --- | --- | --- | --- | --- | --- | --- | --- | --- | --- | --- |
|  |  |  |  |  |  | minR<br>10 | minR<br>5 | minR<br>3 | minR<br>10 | minR<br>5 | minR<br>3 |
| N8 | 9,256 | 1,285,218,728 | 138,852 | 5,559 | 9,115 | 366 | 1,567 | 7,253 | 47 | 181 | 552 |
| N7 | 8,478 | 1,579,342,505 | 186,287 | 6,049 | 8,125 | 431 | 1,994 | 9,032 | 57 | 184 | 568 |
| N5 | 8,906 | 1,071,156,174 | 120,273 | 5,439 | 8,533 | 277 | 1,134 | 5,357 | 23 | 134 | 422 |

### 3.2. Position-based SNVs annotation

To assess the distribution of the SNVs in regard to their position in the gene and predicted functionality, we annotated the SNVs via SeattleSeq (v13, dbSNP build 150); the distribution of functional annotations at each of the three thresholds is shown on Figure 3.

**Figure 3.**
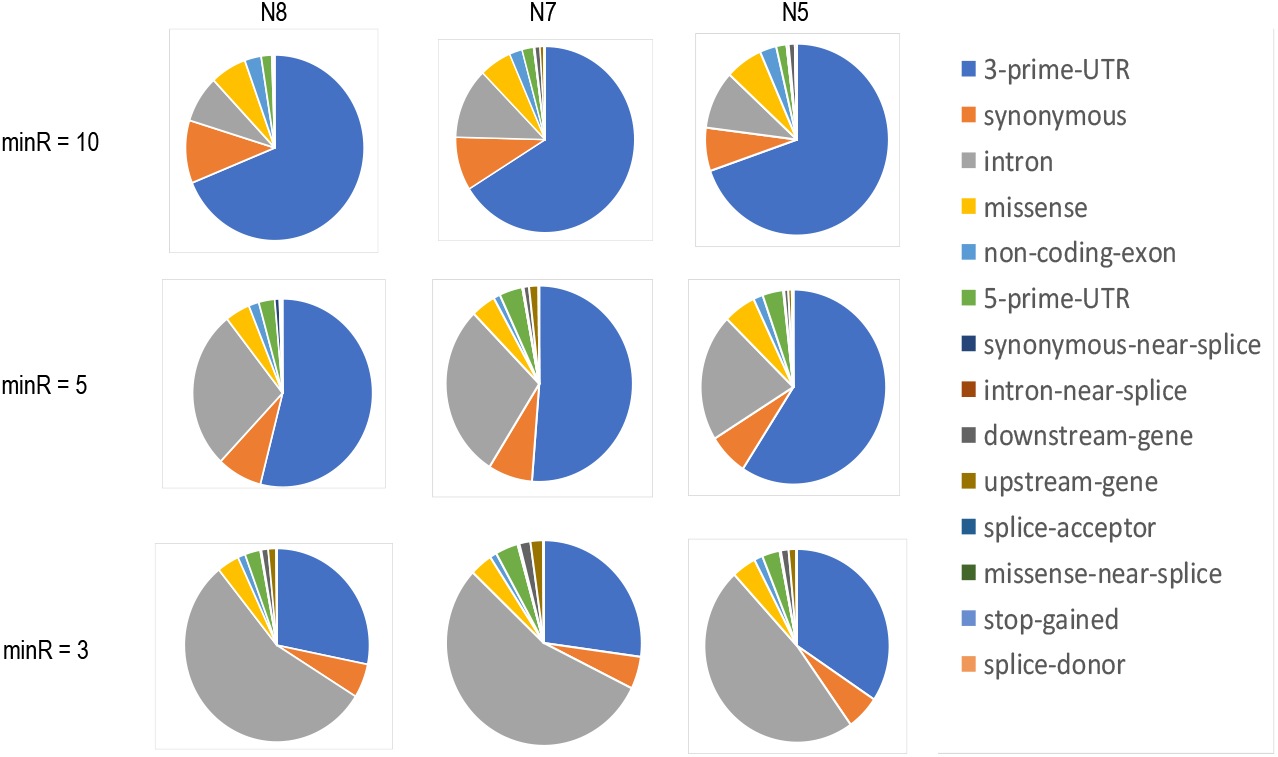
Functional annotation (based on the position in regard to the harboring genes) of SNVs captured by the 10x Genomics platform with different required minimal count of unique sequencing reads. At minR = 5, over 45% of the SNVs are positioned downstream from the 3’-UTR regions.

At minR = 10, close to three-quarters of the captured SNV loci were positioned in the 3’-UTRs of the transcripts, while at minR = 5 this proportion decreased to slightly over 50%. At minR = 3 approximately a quarter of the captured SNVs were positioned in the 3’UTR, while the intronic SNVs increased in proportion to more than 50%. At all thresholds, over 15% of the SNVs were exonic. The complete annotations are shown in Supplementary Tables 1-3.

### 3.3. Allelic expression from single cells at SNV level

To assess allelic expression from single cells, we analyzed VAF_RNA_ at all SNV loci covered with the required number of sequencing reads (minR = 10, 5 and 3), in at least 10 individual cells. For each VAF_RNA_ estimation in each cell, we computed a number of statistics, including mean, median, and percentage of mono- and bi-allelic expressing cells (See also Supplementary Tables 1-3). At all thresholds, the distribution of the VAF_RNA_ mean and median values was generally symmetrical in regard to the VAF_RNA_ scale (Supplementary Figure 1). At all thresholds, more than half of the SNVs presented with bi-allelic expression (0.2<VAF_RNA_<0.8, Table 2). Specifically, VAF_RNA_ obtained at minR = 3 shows biallelic expression for over 50% of the SNVs, and this proportion increases to approximately 90% when confining the analysis to VAF_RNA_ estimated at minR = 10.

**Table 2.** Percent mono- and bi-allelic expression of SNVs covered with different required minimum count od sequencing reads. *\*Predominantly monoallelic expression is inclusive for strict monoallelic expression*.

| Sample | % strictly monoallelic<br>$VAF_{RNA} = 0 \text{ or } 1$ | | | % predominantly monoallelic*<br>$VAF_{RNA} = 0 - 0.2 \text{ or } 0.8 - 1$ | | | % biallelic<br>$VAF_{RNA} = 0.2 - 0.8$ | | |
| --- | --- | --- | --- | --- | --- | --- | --- | --- | --- |
|  | minR 10 | minR 5 | minR 3 | minR 10 | minR 5 | minR 3 | minR 10 | minR 5 | minR 3 |
| N8 | 4.4 | 15.1 | 38.2 | 16.1 | 30.6 | 45.5 | 83.9 | 69.4 | 54.5 |
| N7 | 2.9 | 15.8 | 41.3 | 13.1 | 30.3 | 47.9 | 86.9 | 69.7 | 52.1 |
| N5 | 2.4 | 12.5 | 35.9 | 10 | 26.1 | 42.0 | 90 | 76.9 | 58.0 |

The distribution of scVAF_RNA_ estimations at minR = 10, 5 and 3 for all the heterozygous SNVs in the corresponding sample is shown on Figure 4; the histograms include bins for strictly mono-allelic expression, defined as VAF_RNA_ values of 0 and 1.

**Figure 4.**
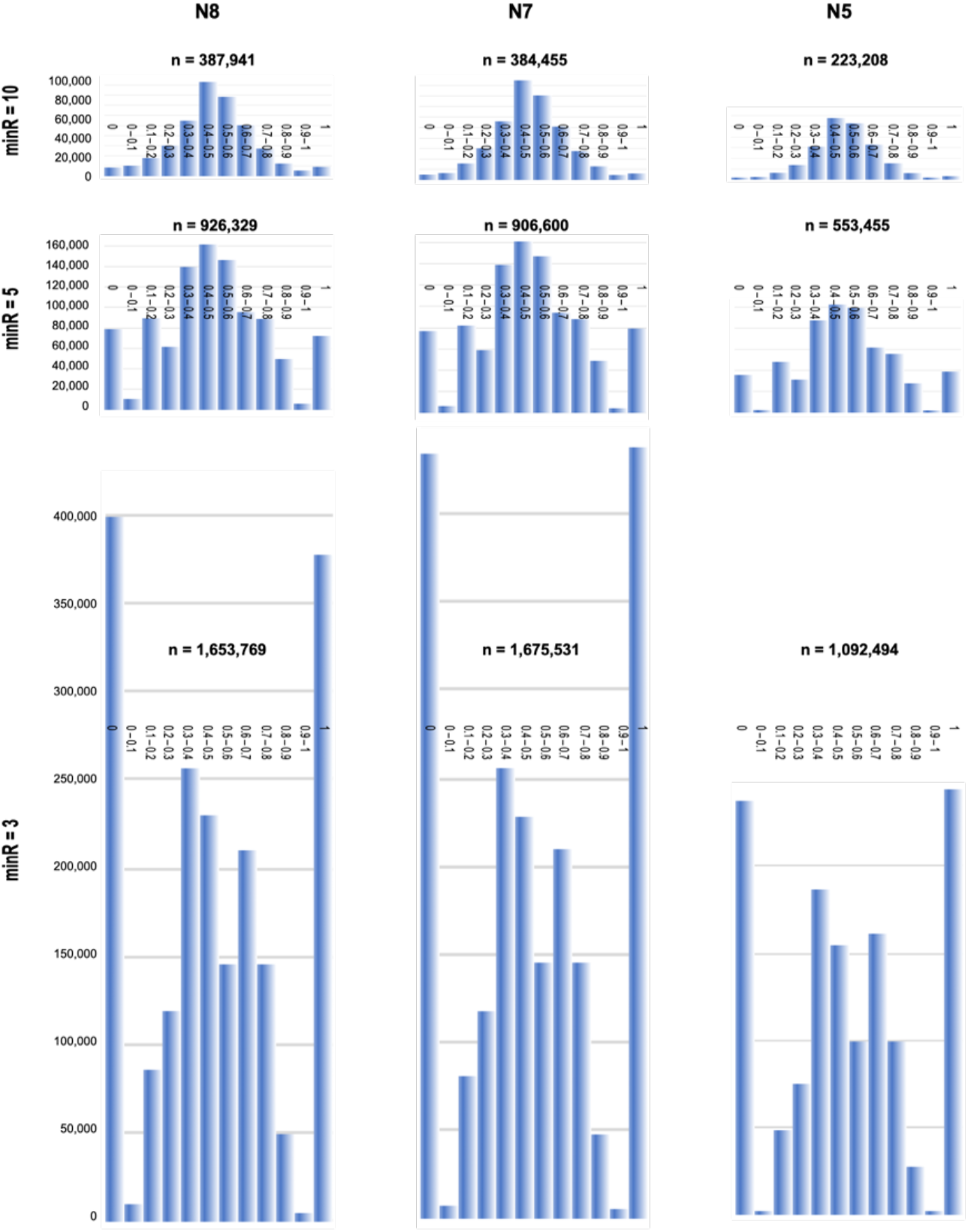
Histograms representing the distribution of scVAF_RNA_ at minR = 10 (top), minR =5 (middle) and minR = 3 (bottom) for all the heterozygote SNVs in N8, N7 and N5. The bin width (x-axes) is 0.1; bin intervals are indicated in the middle of each plot. The y-axes show the numbers of VAF_RNA_ measurements in the individual cells. The total number of VAF_RNA_ estimations (n, across all the cells per group) is shown on the top of each histogram. The histograms are scaled in regards to number of cells. Across the entire dataset, at minR=10 and minR=5, the majority of the SNVs showed biallelic expression centered around VAF_RNA_ of 0.5 (0.4 < VAF_RNA_ < 0.6). In contrast, at minR = 3, the majority of the SNVs presented with strict monoallelic expression (VAF_RNA_ = 0 or 1). The VAFRNAdistributions showed remarkable similarity across the three individuals (N8, N7 and N5).

We next compared our observations to one of the largest previous studies that uses VAF_RNA_ on human scRNA-seq data [17]. The VAF_RNA_ distribution in our data observed at minR = 3 is similar to that in Borel *et al* [17] and is consistent with frequent RME of low- and moderately expressed genes [14–17]. In contrast, at minR = 5, strictly mono-allelic VAF_RNA_ measurements represented less then half of those with VAF_RNA_ = 0.5 ± 0.1, while at minR = 10 we observed gradual decrease of the number of measurements from VAF_RNA_ = 0.5 ± 0.1, towards VAF_RNA_ values of 0 or 1.

Next, we analyzed the data per SNV, across all the cells from which the VAF_RNA_ was estimated at the required minR. For this analysis, to compare findings to Borel *et al* [17], we used similar definitions for allelic expression. Specifically, as monoallelic expression (including RME) we defined SNVs for which fewer than 5% of the cells displayed VAF_RNA_ between 0.2 and 0.8 (0.2<VAF_RNA_<0.8). Skewed allelic expression was assigned to SNVs where less than 10% of the cells expressed one type of allele and the rest expressed either the second allele or both alleles (<80% cells with 0.2 < VAF_RNA_ < 0.8).

VAF_RNA_ distributions for all the SNVs (genome-wide) assessed from a minimum of 1000 cells are plotted on Figure 5. Aligned with the above observations, at minR = 10, the majority of the SNV loci, had biallelic expression, with substantial proportion of the cells having scVAF_RNA_ estimations between 0.2 and 0.8; this proportion gradually decreased at the lower thresholds, but remained above 50% at minR = 3.

**Figure 5.**
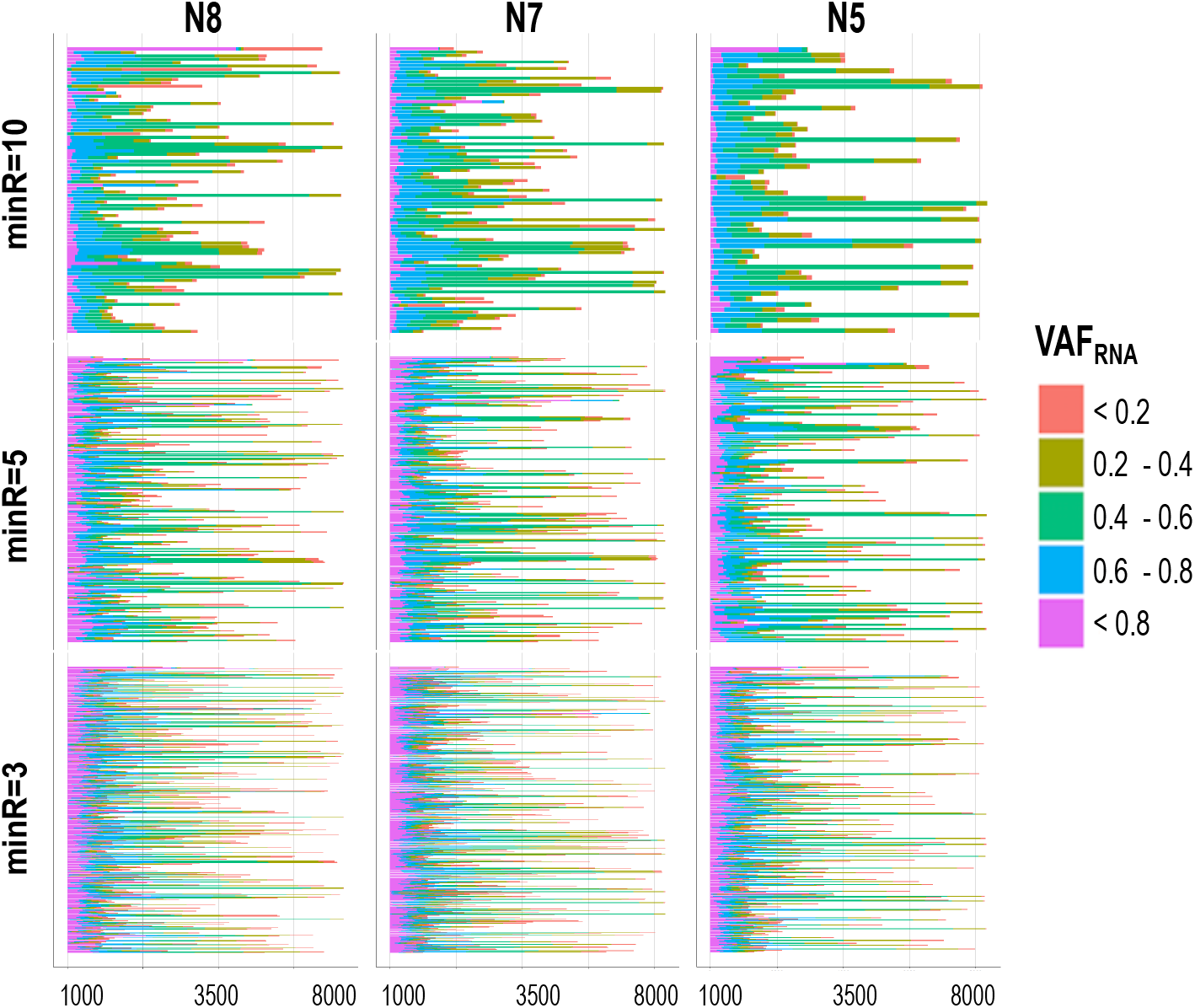
scVAF_RNA_ estimated at positions covered by a minimum of 10 sequencing reads (top), 5 sequencing reads (middle) and 3 sequencing reads (bottom) across more than 1000 cells. For the majority of the positions, VAF_RNA_ showed biallelic expression, with substantial proportion of the scVAF_RNA_ estimations in the interval 0.4 – 0.6.

### 3.4. Allelic expression from single cells at gene level

We analyzed allele-specific expression at gene level, again, comparing our findings to [29]. We first performed this analysis at minR=10, at which cutoff 21 genes from our dataset overlapped with the 60 genes highlighted in Borel *et al* [17] (Figure 6, top); all 21 genes showed biallelic expression in a complete agreement. From the above-mentioned 60 genes, autosomal genes with RME were observed only at minR = 3 in our dataset, all of them in complete concordance with Borel *at el* [17]. Examples of such genes are shown (Figure 6, bottom), including the strictly monoallelic *RAD52*. Out of the 12 genes with reported skewed allelic expression, 4 were present in our dataset: *CNN3*, *C12orf75* and *CCDC80* had skewed expression, while *SPC3* showed symmetrically distributed alleles in both samples where it was detected (See Supplementary Table 3).

**Figure 6.**
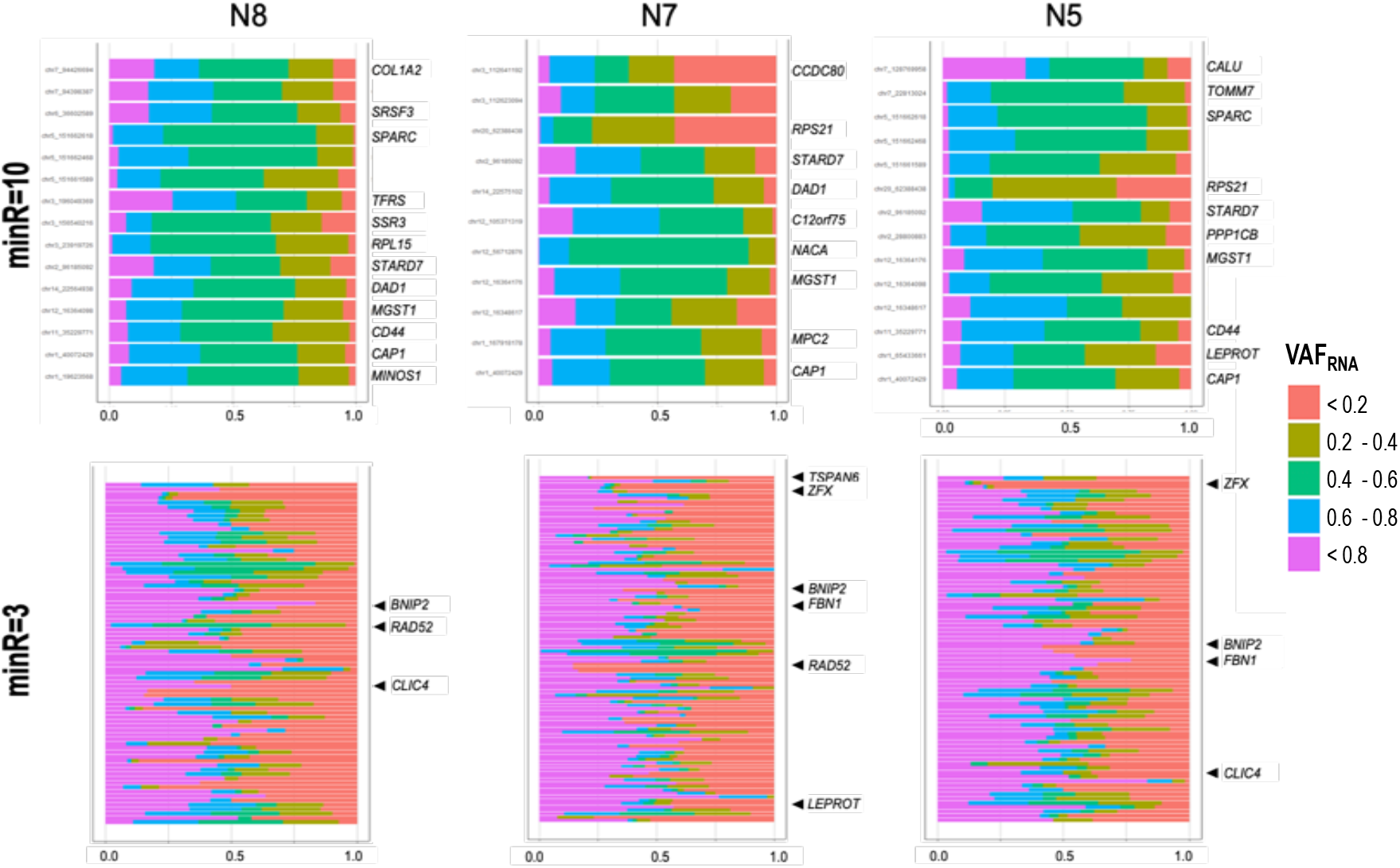
scVAF_RNA_ distribution at positions covered by a minimum of 10 sequencing reads (top), and 3 sequencing reads (bottom) across more than 1500 cells for genes reported by Borel *et al* [17]. For the positions with minR = 10, no RME was suggested by the scVAF_RNA_ distribution for autosomal genes (i.e. multiple scVAF_RNA_ values between 0.4 – 0.6, while positions covered with minR = 3 showed frequent monoallelic signals (scVAF_RNA_ > 0.8 or scVAF_RNA_ < 0.2. As expected, chrX shows strong RME patterns (see gene *TSPAN6*).

In this gene-set, for genes with multiple SNVs, we observed concordant allelic expression (See *COL1A2*, *SPARC*, *CCDC80*, and *MGST1* on Figure 6). We also observed complete concordance across the three individuals for the SNVs shared between donors; SNVs common for the three donors and assessed from more than 50 cells per donor are shown on Figure 7 (chromosome 1, the rest of the chromosomes showed similar results; see also *CAP1*, *DAD1*, *SPARC*, *MGST1*, *CD44*, *STARD7* on Figure 6).

**Figure 7.**
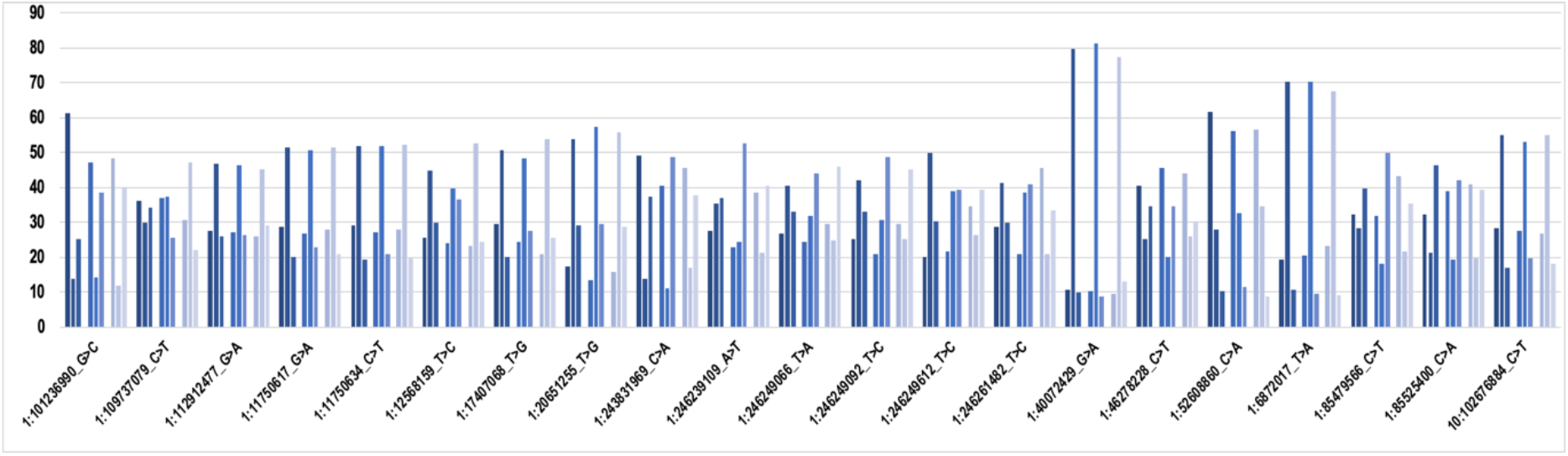
Percentage of cells (y-axis) displaying VAF_RNA_<0.2 (for each cluster of three, left), VAF_RNA_ between 0.2 and 0.8 (middle), and VAF_RNA_ > 0.8 (right) in the three donors (N5 – dark blue, N7-light blue, N8 – grey). High concordance between the three donors is seen; SNVs on chromosome 1 are shown, the results were similar genome-wide.

Next, we assessed mono- and bi-allelic SNV-expression at gene level across our entire dataset. We confined this assessment to SNVs seen in a minimum of 50 cells per sample; 7408 SNVs in 3406 genes were eligible for this analysis at minR=3 across the three donors. Predominant RME (fewer than 5% of the cells with VAF_RNA_ between 0.2 and 0.8) was seen in 451 SNVs positioned in 376 genes; from those, 49 SNVs in 42 genes did not have any cells expressing both alleles.

We next assessed if the allelic status is consistent across multiple SNVs from the same gene. To do this, we pooled the SNVs from the three donors together and selected genes with more than 3 SNVs, each assessed from a minimum of 50 cells per donor; 3922 SNVs in 815 genes were available for this analysis. The first striking observation was that, for the majority of the genes, intronic SNVs have substantially higher rates of monoallelic expression, as compared to SNVs in the spliced mRNA (Figure 8). This was evident both at the level of the individual genes (examples shown on Figure 8a), and genome-wide, where the average proportion of cells expressing both alleles (0.2<VAF_RNA_< 0.8) was significantly lower for SNV positioned in introns, as compared to SNVs in exons and UTRs (Figure 8b). Aligned with the above, the proportion intronic SNVs was significantly higher among the genes with RME (p < 2.2e-16, chi-square test, Figure 8c). Within the groups of intronic and non-intronic SNVs in the same gene, highly consistent VAF_RNA_ distributions were observed.

**Figure 8.**
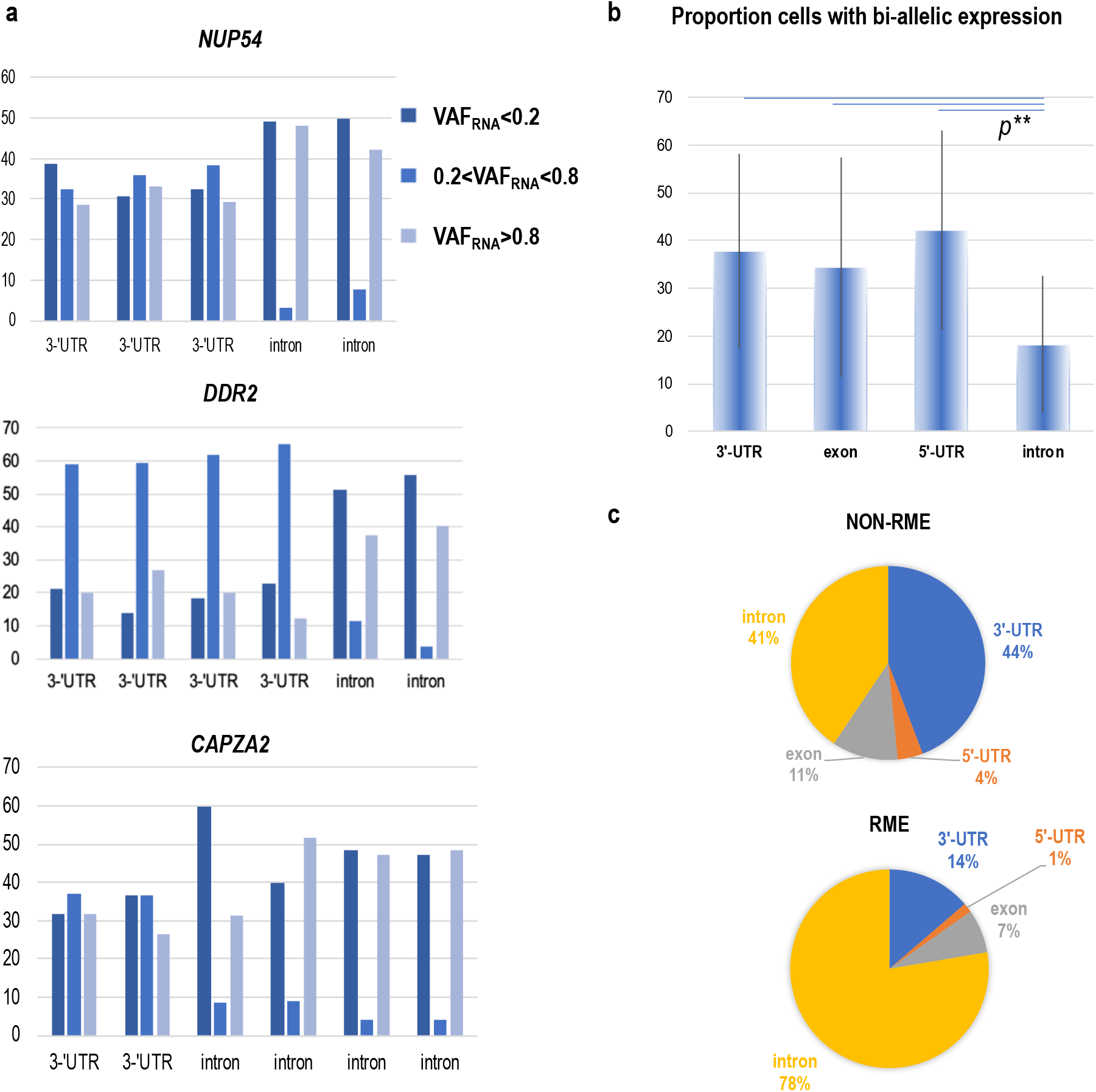
**a.** Genes with multiple SNVs positioned in intronic and non-intronic sequences; high percentage of RME cells (VAF_RNA_<0.2 or >0.8, y-axis) is obvious. **b**. Average percentage of cells (y-axis) with biallelic expression across all SNVs in our dataset stratified by position in the gene; SNVs positioned in introns were be-allelic in a lower proportion of cells as compared to all other SNVs. **c**. Distribution of position annotations in Non-RME and RME groups of SNVs; substantially higher proportion of intronic loci is seen among the RME-SNVs.

## 4. Discussion

Our analysis includes more than 4 billion RNA-seq reads and over 7.8 million individual scVAF_RNA_ estimations, making it, to our knowledge, the largest study on SNV-based allele-specific expression from human scRNA-seq data. We note that, as we do not have the genotypes to select heterozygous loci, we confined our analysis to SNVs that are confidently bi-allelic (a minimum of 50 reads supporting each allele from the pooled RNA-seq data per donor). By default, this selection excludes heterozygous SNVs with strong non-random mono-allelic expression (which would appear as monoallelic in the pooled RNA-seq data). Therefore, the presented here results need to be considered strictly in the light of this selection.

We use the advantage of the large number of cells (over 24K) and high sequencing depth (150K reads per cell) to explore the feasibility of scVAF_RNA_ estimations, and define a set of scVAF_RNA_ characteristics. Our results show that SNV assessment from scRNA-seq generated through the 3’-based 10x Genomics platform can be highly informative for several reasons.

First, the annotation of the captured variants (See Figure 3) supports analyses on variant functionality. As expected, 10x-Genomics scRNA-seq data contains a significant proportion of the gene 3’-UTR variants, which are known to strongly affect both gene expression and splicing [37–41]. In addition, approximately 15% of the captured SNVs are exonic, and include missense, nonsense, and near-splice variants, many of which can potentially affect the protein structure and function (See Supplementary Tables 1-3). Importantly, the platform captures a substantial number of intronic SNVs. Intronic sequences are reported in 15-25% of the RNA-sequencing reads from both bulk and single-cell based studies [18, 42, 43]. ScRNA-seq intronic sequences can be used to estimate the relative abundance of precursor and mature mRNA, thereby assessing the RNA velocity, and respectively, cellular dynamic processes [18]. Consistent with a major recent study on RNA velocity [18] and models of transcriptional burst kinetics [13–15] we observe significantly higher monoallelic expression for intronic SNVs as compared to non-intronic SNVs in the same gene (See Figure 8). Specifically, it is established that at times of increased transcription, unspliced precursors are rapidly produced (often from one of the alleles), and conversely the proportion of unspliced mRNAs is quickly lessened during times of reduced transcription. Therefore, at any given moment a single cell is likely to contain unspliced precursors produced from one of the alleles, compared to the longer-living spliced mRNAs of the same gene, which are more likely to accumulate both alleles over time. Because the balance of unspliced and spliced mRNA abundance is predictive for the future state of the mature RNA [18], scVAF_RNA_ analyses can be applied to assess dynamic cellular processes.

Second, to our knowledge, this is the first study to estimate allele expression from a minimum of 10 unique sequencing reads from scRNA-seq data. Our findings mostly agree with previous studies and also indicate, that at such stringency the majority of the autosomal genes show largely symmetric biallelic expression. This effect is expected given that genes captured with a minimum of 10 unique sequencing reads from a single cell correspond to highly expressed genes, which are frequently associated with house-keeping functions [44]. We provide this data (minR = 10, Supplementary Table 1), together with the estimations at minR = 5 and minR = 3 (Supplementary Tables 2 and 3) which can be used for analyses of allele-specific expression both genome-wide and at the level of individual genes of interest.

Third, we present a set of characteristics of VAF_RNA_ obtained from scRNA-seq data. Several factors facilitate the applicability of VAF_RNA_ to assess functional genetic variants (from both bulk and scRNA-seq data). First, as mentioned earlier, VAF_RNA_ allows for precise allele quantitation, particularly important for sites with allele-specific regulation, RNA-editing, and somatic mutations in cancer. Here it is important to note that allele mapping bias might be critical for accurate VAF_RNA_ estimation, especially from loci with a low number of sequencing reads as is common in scRNA-seq data. We correct our alignments for allele-mapping bias using the WASP package [32]. WASP is implemented in the latest versions of the herein used popular alignment STAR [20], which significantly streamlines the data processing, especially for datasets with predefined lists of SNV loci of interest (i.e. available genotypes, lists of known SNVs of interest such as RNA-editing, dbSNP, etc.). Second, VAF_RNA_ is dynamic and reflects the actual allele content in the cell at a particular moment in time. In scRNA studies, where the different cells are commonly in gradual states of progressive processes, VAF_RNA_ analyses can be adopted to study lineages and cellular dynamics. Third, VAF_RNA_ can be used to study functional SNVs from sets where matched DNA (and, respectively genotypes) is not available [24,25]. Ultimately, these analyses apply to expressed SNVs and will not capture loci positioned in transcriptionally silent regions. The single-cell resolution of this approach brings further advantages. First, due to preservation of intracellular relationships between molecular features, single-cell analyses facilitate the discovery of correlations between SNVs and other transcriptome features, such as gene expression or splicing. Finally, scRNA-seq projects typically utilize cells with (largely) identical genotypes (i.e. from the same system/individual), thus supplying context for assessment of SNVs implicated in RNA-specific regulation.

## 5. Conclusions

In conclusion, we present a large SNV-focused study on allele expression from scRNA-seq data that addresses three major technical factors known to bias single cell allelic studies: PCR-related bias, allele-mapping bias, and a low number of sequencing reads. To facilitate similar studies, we describe a step-by-step approach for confident scVAF_RNA_ estimations. Our study is largely consistent with existing knowledge, reports findings on previously unassessed genes and SNVs, and supplies datasets for further analyses. In addition, our analysis demonstrates the feasibility of scVAF_RNA_ estimation from current scRNA-seq datasets and shows that the 3’-based library generation protocol of 10x Genomics scRNA-seq data can be highly informative in SNV-based analyses.

## Supplementary Materials

The following are available online at www.mdpi.com/xxx/s1, S_Figure_1_Mean_and_Median_VAF_RNA_, S_Table_1_SNV_loci_minR10_10cells, S_Table_2_SNV_loci_minR5_10cells, S_Table_3_SNV_loci_minR10_10cells.

## Author Contributions

Conceptualization, writing—original draft preparation, and supervision, A.H; methodology, software, visualization and writing—review and editing, NMP, HL, PB, LS, NA, HI, JS, DRS.

## References

1. Kulkarni A, Anderson AG, Merullo DP, Konopka G. Beyond bulk: a review of single cell transcriptomics methodologies and applications. Curr Opin Biotechnol 2019, 58, 129–136

2. Stuart T, Satija R. Integrative single-cell analysis. Nat Rev Genet 2019, 20, 257–272.

3. Zafar H, Wang Y, Nakhleh L, Navin N, Chen K. Monovar: single-nucleotide variant detection in single cells. Nat Methods 2016, 13, 505–507.

4. Schnepp PM, Chen M, Keller ET, Zhou X. SNV identification from single-cell RNA sequencing data. Hum Mol Genet 2019.

5. Dong M, Jiang Y. Single-Cell Allele-Specific Gene Expression Analysis. Methods Mol Biol 2019, 1935, 155–174.

6. Griffiths JA, Scialdone A, Marioni JC. Mosaic autosomal aneuploidies are detectable from single-cell RNAseq data. BMC Genomics 2017, 18, 904.

7. Moreira de Mello JC, Fernandes GR, Vibranovski MD, Pereira LV. Early X chromosome inactivation during human preimplantation development revealed by single-cell RNA-sequencing. Sci Rep 2017, 7, 10794.

8. Edsgärd D, Reinius B, Sandberg R. scphaser: haplotype inference using single-cell RNA-seq data. Bioinformatics 2016, 32, 3038–3040.

9. Kim JK, Kolodziejczyk AA, Ilicic T, Teichmann SA, Marioni JC. Characterizing noise structure in single-cell RNA-seq distinguishes genuine from technical stochastic allelic expression. Nat Commun 2015, 6, 8687.

10. Poirion O, Zhu X, Ching T, Garmire LX. Using single nucleotide variations in single-cell RNA-seq to identify subpopulations and genotype-phenotype linkage. Nat Commun 2018, 9, 4892.

11. Vu TN, Nguyen HN, Calza S, Kalari KR, Wang L, Pawitan Y. Cell-level somatic mutation detection from single-cell RNA-sequencing. Bioinformatics 2019.

12. Rodriguez-Meira A, Buck G, Clark SA, Povinelli BJ, Alcolea V, Louka E, McGowan S, Hamblin A, Sousos N, Barkas N, Giustacchini A, Psaila B, Jacobsen SEW, Thongjuea S, Mead AJ. Unravelling Intratumoral Heterogeneity through High-Sensitivity Single-Cell Mutational Analysis and Parallel RNA Sequencing. Mol Cell 2019, 73, 1292–1305.e8.

13. Larsson AJM, Johnsson P, Hagemann-Jensen M, Hartmanis L, Faridani OR, Reinius B, Segerstolpe Å, Rivera CM, Ren B, Sandberg R. Genomic encoding of transcriptional burst kinetics. Nature 2019, 565, 251–254.

14. Kim JK, Marioni JC. Inferring the kinetics of stochastic gene expression from single-cell RNA-sequencing data. Genome Biol 2013, 14, R7.

15. Deng Q, Ramsköld D, Reinius B, Sandberg R. Single-cell RNA-seq reveals dynamic, random monoallelic gene expression in mammalian cells. Science 2014, 343, 193–196.

16. Reinius B, Mold JE, Ramsköld D, Deng Q, Johnsson P, Michaëlsson J, Frisén J, Sandberg R. Analysis of allelic expression patterns in clonal somatic cells by single-cell RNA-seq. Nat Genet 2016, 48, 1430–1435.

17. Borel C, Ferreira PG, Santoni F, Delaneau O, Fort A, Popadin KY, Garieri M, Falconnet E, Ribaux P, Guipponi M, Padioleau I, Carninci P, Dermitzakis ET, Antonarakis SE. Biased allelic expression in human primary fibroblast single cells. Am J Hum Genet 2015, 96, 70–80.

18. La Manno G, Soldatov R, Zeisel A, Braun E, Hochgerner H, Petukhov V, Lidschreiber K, Kastriti ME, Lönnerberg P, Furlan A, Fan J, Borm LE, Liu Z, van Bruggen D, Guo J, He X, Barker R, Sundström E, Castelo-Branco G, Cramer P, Adameyko I, Linnarsson S, Kharchenko PV. RNA velocity of single cells. Nature 2018, 560, 494–498.

19. Horvath A, Pakala SB, Mudvari P, Reddy SD, Ohshiro K, Casimiro S, Pires R, Fuqua SA, Toi M, Costa L, Nair SS, Sukumar S, Kumar R. Novel insights into breast cancer genetic variance through RNA sequencing. Sci Rep 2013, 3, 2256.

20. Van der Auwera GA, Carneiro MO, Hartl C, Poplin R, Del Angel G, Levy-Moonshine A, Jordan T, Shakir K, Roazen D, Thibault J, Banks E, Garimella KV, Altshuler D, Gabriel S, DePristo MA. From FastQ data to high confidence variant calls: the Genome Analysis Toolkit best practices pipeline. Curr Protoc Bioinformatics 2013, 43.

21. Deelen P, Zhernakova DV, de Haan M, van der Sijde M, Bonder MJ, Karjalainen J, van der Velde KJ, Abbott KM, Fu J, Wijmenga C, Sinke RJ, Swertz MA, Franke L. Calling genotypes from public RNA-sequencing data enables identification of genetic variants that affect gene-expression levels. Genome Med 2015, 7, 30.

22. Kravitz SN, Gregg C. New subtypes of allele-specific epigenetic effects: implications for brain development, function and disease. Curr Opin Neurobiol 2019, 59, 69–78.

23. Lee MP. Understanding Cancer Through the Lens of Epigenetic Inheritance, Allele-Specific Gene Expression, and High-Throughput Technology. Front Oncol 2019, 9, 794.

24. Spurr L, Alomran N, Bousounis P, Reece-Stremtan D, Prashant NM, Liu H, Słowiński P, Li M, Zhang Q, Sein J, Asher G, Crandall KA, Tsaneva-Atanasova K, Horvath A. ReQTL: Identifying correlations between expressed SNVs and gene expression using RNA-sequencing data. Bioinformatics 2019.

25. Sein J, Spurr L, Bousounis P, Prashant NM, Liu H, Alomran N, Bernot J, Ibeawuchi H, Reece-Stremtan D, Horvath A. RsQTL: correlation of expressed SNVs with splicing using RNA-sequencing data. Bioinformatics 2019. Under Review https://www.biorxiv.org/content/10.1101/840504v1.

26. Movassagh M, Alomran N, Mudvari P, Dede M, Dede C, Kowsari K, Restrepo P, Cauley E, Bahl S, Li M, Waterhouse W, Tsaneva-Atanasova K, Edwards N, Horvath A. RNA2DNAlign: nucleotide resolution allele asymmetries through quantitative assessment of RNA and DNA paired sequencing data. Nucleic Acids Res 2016; 44, e161.

27. Mudvari P, Movassagh M, Kowsari K, Seyfi A, Kokkinaki M, Edwards NJ, Golestaneh N, Horvath A. SNPlice: variants that modulate Intron retention from RNA-sequencing data. Bioinformatics 2015, 31, 1191–1198.

28. Restrepo P, Movassagh M, Alomran N, Miller C, Li M, Trenkov C, Manchev Y, Bahl S, Warnken S, Spurr L, Apanasovich T, Crandall K, Edwards N, Horvath A. Overexpressed somatic alleles are enriched in functional elements in Breast Cancer. Sci Rep 2017, 7, 8287.

29. Spurr L, Li M, Alomran N, Zhang Q, Restrepo P, Movassagh M, Trenkov C, Tunnessen N, Apanasovich T, Crandall KA, Edwards N, Horvath A. Systematic pan-cancer analysis of somatic allele frequency. Sci Rep 2018, 8, 7735.

30. Suvà ML, Tirosh I.Single-Cell RNA Sequencing in Cancer: Lessons Learned and Emerging Challenges. Mol Cell 2019, 75, 7–12.

31. Liu X, Xiang Q, Xu F, Huang J, Yu N, Zhang Q, Long X, Zhou Z. Single-cell RNA-seq of cultured human adipose-derived mesenchymal stem cells. Sci Data 2019, 6, 190031.

32. van de Geijn B, McVicker G, Gilad Y, Pritchard JK. WASP: allele-specific software for robust molecular quantitative trait locus discovery. Nat Methods 2015, 12, 1061–3.

33. Smith T, Heger A, Sudbery I. UMI-tools: modeling sequencing errors in Unique Molecular Identifiers to improve quantification accuracy. Genome Res 2017, 27, 491–499.

34. Dobin A, Davis CA, Schlesinger F, Drenkow J, Zaleski C, Jha S, Batut P, Chaisson M, Gingeras TR. STAR: ultrafast universal RNA-seq aligner. Bioinformatics 2013, 29, 15–21.

35. Liao Y, Smyth GK, Shi W. featureCounts: an efficient general purpose program for assigning sequence reads to genomic features. Bioinformatics 2014, 30, 923–930.

36. Butler A, Hoffman P, Smibert P, Papalexi E, Satija R. Integrating single-cell transcriptomic data across different conditions, technologies, and species. Nat Biotechnol 2018, 36, 411–420.

37. Gruber AJ, Gypas F, Riba A, Schmidt R, Zavolan M. Terminal exon characterization with TECtool reveals an abundance of cell-specific isoforms. Nat Methods 2018, 15, 832–836.

38. Kishore S, Luber S, Zavolan M. Deciphering the role of RNA-binding proteins in the post-transcriptional control of gene expression. Brief Funct Genomics 2010, 9, 391–404.

39. Hausser J, Zavolan M. Identification and consequences of miRNA-target interactions--beyond repression of gene expression. Nat Rev Genet 2014, 15, 599–612.

40. Chatterjee S, Pal JK. Role of 5’-and 3’-untranslated regions of mRNAs in human diseases. Biol Cell 2009, 101, 251–62.

41. Maiti GP, Ghosh A, Mondal P, Baral A, Datta S, Samadder S, Nayak SP, Chakrabarti J, Biswas J, Sikdar N, Chowdhury S, Roy B, Roychowdhury S, Panda CK. SNP rs1049430 in the 3’-UTR of SH3GL2 regulates its expression: Clinical and prognostic implications in head and neck squamous cell carcinoma. Biochim Biophys Acta 2015, 1852, 1059–1067.

42. Picelli S, Björklund ÅK, Faridani OR, Sagasser S, Winberg G, Sandberg R. Smart-seq2 for sensitive full-length transcriptome profiling in single cells. Nat Methods 2013, 10, 1096–8.

43. Gaidatzis D, Burger L, Florescu M, Stadler MB.Analysis of intronic and exonic reads in RNA-seq data characterizes transcriptional and post-transcriptional regulation. Nat Biotechnol 2015, 33, 722–729.

44. Tani H, Mizutani R, Salam KA, Tano K, Ijiri K, Wakamatsu A, Isogai T, Suzuki Y, Akimitsu N. Genome-wide determination of RNA stability reveals hundreds of short-lived noncoding transcripts in mammals. Genome Res 2012, 22, 947–56.

