## Supplementary Tables 1-3 for "Estimating allele-specific expression of SNVs from 10x Genomics Single-Cell RNA-Sequencing Data"

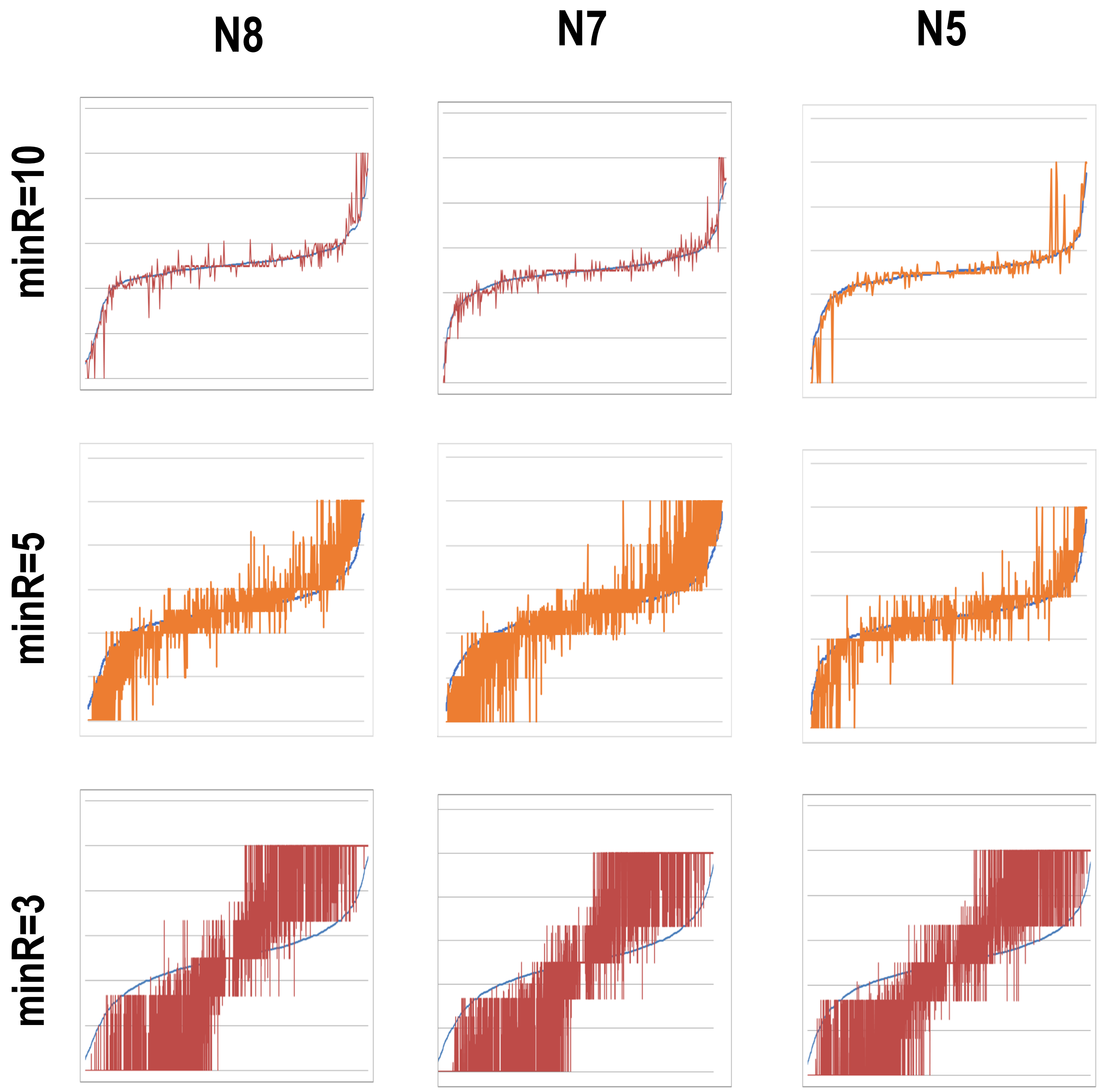


Supplementary Figure 1: Mean (blue) and Median (orange) $VAF_{RNA}$ at different thresholds of required minimum sequencing reads (minR = 10 (top), minR = 5 (middle) and minR = 3 (bottom)/
